# Modeling human eye movements during immersive visual search

**DOI:** 10.1101/2022.12.01.518717

**Authors:** Angela Radulescu, Bas van Opheusden, Frederick Callaway, Thomas L. Griffiths, James M. Hillis

## Abstract

The nature of eye movements during visual search has been widely studied in psychology and neuroscience. Virtual reality (VR) paradigms provide an opportunity to test whether computational models of search can predict naturalistic search behavior. However, existing ideal observer models are constrained by strong assumptions about the structure of the world, rendering them impractical for modeling the complexity of environments that can be studied in VR. To address these limitations, we frame naturalistic visual search as a problem of allocating limited cognitive resources, formalized as a meta-level Markov decision process (meta-MDP) over a representation of the environment encoded by a deep neural network. We train reinforcement learning agents to solve the meta-MDP, showing that the agents’ optimal policy converges to a classic ideal observer model of search developed for simplified environments. We compare the learned policy with human gaze data from a visual search experiment conducted in VR, finding a qualitative and quantitative correspondence between model predictions and human behavior. Our results suggest that gaze behavior in naturalistic visual search is consistent with rational allocation of limited cognitive resources.

## Introduction

Developing models of human visual search is a long-standing problem in psychology and neuroscience (1–3). Visual search is a common task in which humans look for an item (known as a “target”) in a crowded visual environment that consists of various “distractors.” While visual search tasks differ in how targets and distractors are defined, they all require a searcher to respond to observations from the environment with a sequence of eye movements. For instance, to find our keys in a cluttered room, we have to move our eyes to different regions of the scene to process local details. In addition, we may only represent features of objects (e.g., color and size) that provide information about object identity. Generally, visual search requires selectively sampling information from the environment, under time and resource constraints, in service of a goal.

Visual search has traditionally been studied in behavioral paradigms that approximate various aspects of real-world search using static 2D displays and highly restricted head movement. This research has helped uncover different features that drive eye movements (4–6), and has led to the development of several theoretic accounts of search (7–12). One important insight from this work is that “ideal observer” models, which assume that eye movements are optimally selected to gather information, predict behavioral and neural signatures of information processing in multi-sensory perception tasks (13–19). However, because these ideal observer models have been tailored to specific paradigms with well-defined environment structure, they are typically constrained by statistical assumptions about the visual input (7).

Recent work has begun to leverage virtual reality (VR) to study visual search in naturalistic settings (20–22). This opens up the question of whether ideal observer models can generalize to environments in which humans have to search for suprathreshold, 3D targets in an active-viewing, embodied setting. Instantiating ideal observer models of visual search in VR is challenging, however, because such models do not specify what the relevant features for search should be in the real world, nor do they provide a clear account of what objective the agent should optimize when searching through the space of those features.

In this work, we begin addressing these challenges by framing visual search as a trade-off between gathering information and the cost of the associated computation (23–26). This approach—analyzing cognition as the rational allocation of limited resources, or “resource-rational analysis”— has been used to generate models of a range of cognitive processes, explaining observations from research on judgment (27, 28), decision-making (29, 30), and planning (31). We leverage resource-rational analysis in combination with convolutional neural networks (CNNs) and deep reinforcement learning to specify and train an agent that searches for objects in a naturalistic, high-dimensional visual search task. We compare the behavior of the trained agent to human gaze data in a visual search experiment conducted in VR.

Adopting this perspective affords several advances: by training agents to perform visual search over a representation of the environment produced by a deep neural network, we demonstrate that previously proposed ideal observer models are optimal under more general assumptions that include a formal specification of the structure of naturalistic scenes. We provide evidence that the resource-rational policy predicts human gaze behavior in an active-viewing, immersive setting. And we use this framework to systematically ablate features of the model in order to study which of them guide search in naturalistic contexts. Taken together, our results show that naturalistic visual search can be thought of as a resource-rational sequential decision-making process over a deep representation of the environment.

## Results

To address the question of whether ideal observer models of search generalize to naturalistic environments, we conducted a study of visual search in virtual reality (Fig. 1A and B, Methods A). Human participants were immersed in virtual indoor scenes and interacted with the environment using a handheld controller while their gaze was tracked. When immersed in a scene, the participant’s task was to find a target object among several distractors scattered around the room. Each scene was uniquely defined by the room context (e.g., living room), viewpoint (e.g., above the table), target (e.g., pizza box) and distractor set.

**Fig. 1.**
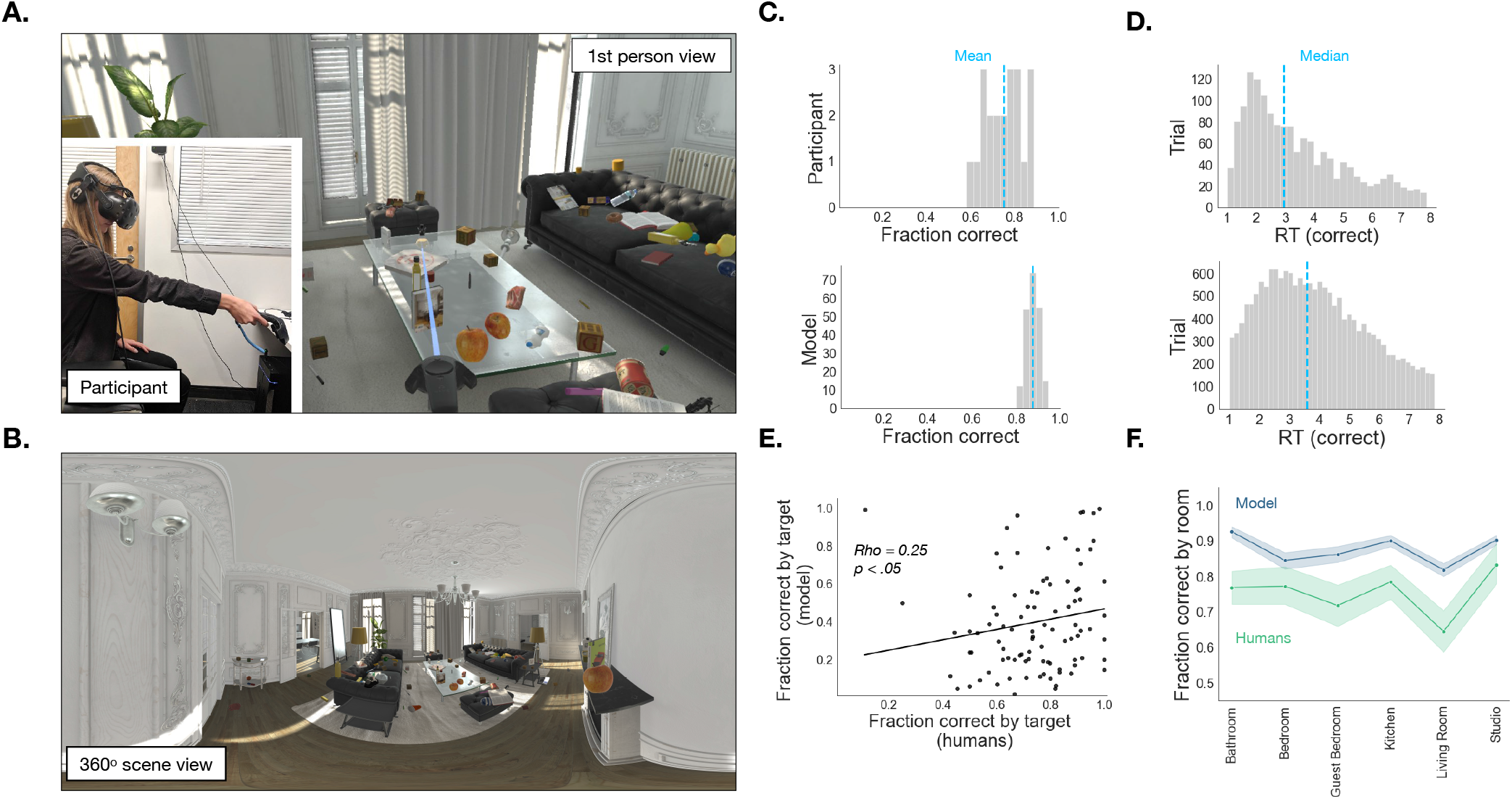
Immersive visual search task. **A**: Egocentric view of the participant shown in the inset during the immersive visual search task. The participant is using the controller to report having found the target (the white pizza box). **B**: 360-degree equirectangular projection of the scene in **A**. Participants experienced the scene from a fixed viewpoint, and were free to move their eyes and head to sample different parts of the environment. **C**: The proportion of times that participants (top) or the model (bottom) correctly identified the target within the allotted 8 seconds across all 300 search episodes. The dashed blue line represents the mean. **D**: Reaction times in all search episodes on which participants (top) or the model (bottom) found the target. The dashed blue line represents the median. **E**: Correlation between actual and simulated average fraction correct as a function of target. **F**: Comparison between actual and simulated average fraction correct as a function of room context. Error bars denote 95% CIs.

Despite the complexity of the environment and the relatively large number of distractors, participants successfully found the target object 75%of the time, fixating an average of 12.6 items before locating the target (Fig. 1C, top). More than half of all successful searches were under 3 seconds long (Fig. 1D, top). Thus, people were able to efficiently search for objects in complex 3D environments. We next asked what strategy they use to accomplish this. In particular, is their fixation behavior consistent with an ideal observer model? To address the question of how people select which object to fixate next, we must first specify how objects are represented.

### A. Defining a feature space

In traditional visual search tasks, simple 2D targets are presented on top of unstructured noisy backgrounds. By contrast, in naturalistic visual search, targets are complex 3D objects that can appear in multiple orientations, scales, and lighting conditions. This necessitates searching for objects that match the target on a set of abstract features that are invariant across pose and illumination—that is, a representation. From a modeling perspective, the choice of representation encodes a formal specification of the prior knowledge that humans bring to the task. To specify this knowledge, we compared two approaches: an explicit approach based on shape and color, and an implicit approach using neural networks.

Previous work in visual cognition has shown that during visual search, eye movements are guided by top-down attention to perceptual features of the target object (5). But which perceptual features do people use? Two likely candidates are shape, quantified using the D2 distribution (32), and color, quantified in the CIE L*a*b* space (Fig. 2A; see Methods B for details). Each of these representations encodes each object as a point in a high-dimensional perceptual space.

**Fig. 2.**
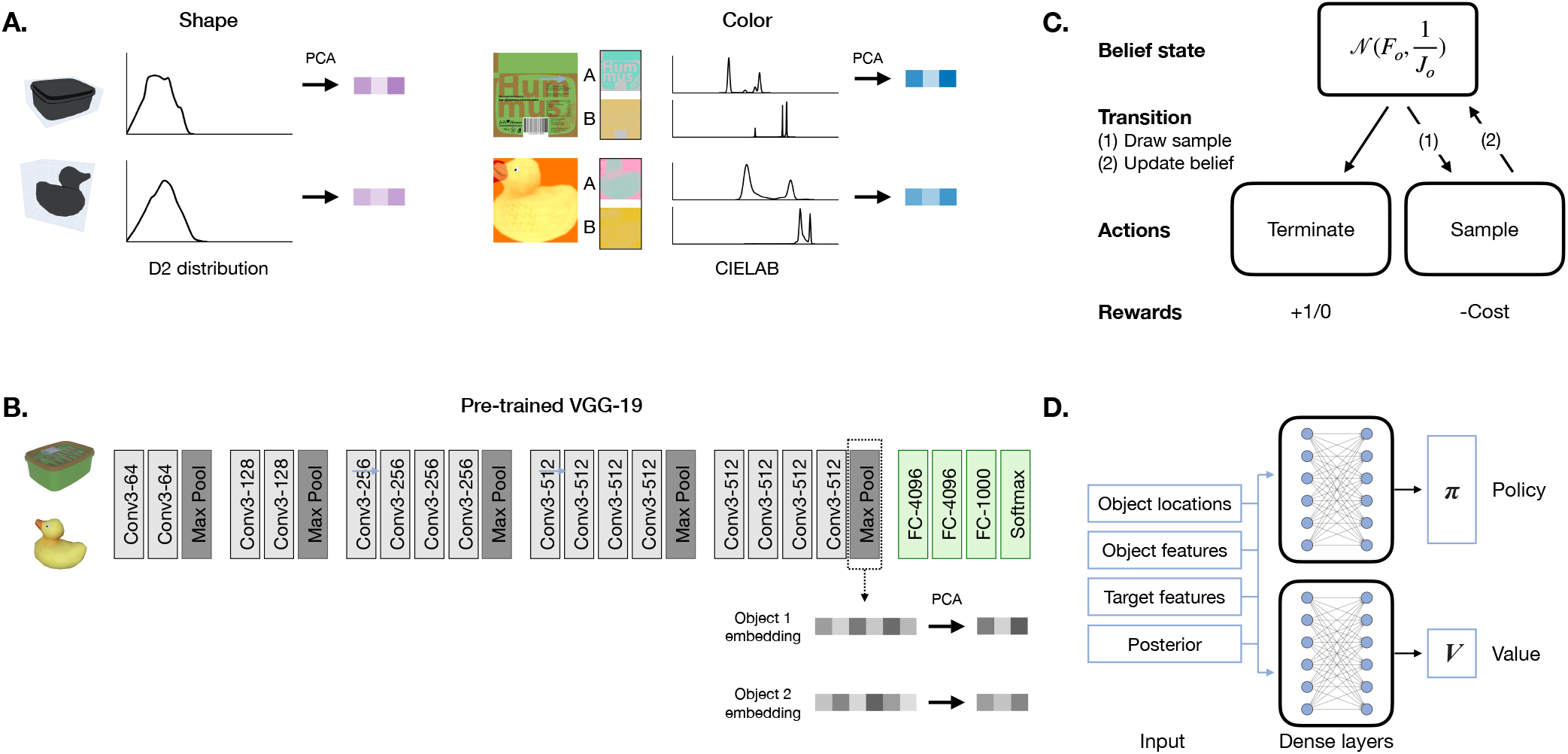
Modeling approach. The feature space of the task consists of objects and their perceptual features. Perceptual features were extracted in two different ways (Methods B): explicitly, by summarizing each object’s shape and color information; and implicitly, by embedding objects in a feature space defined by activation in the layers of a pre-trained convolutional neural network (CNN). **A**: Left, mesh representation and corresponding D2 shape distributions. Right, 2D texture and corresponding CIE L*a*b* color distributions. **B**: Architecture of VGG-19, the pre-trained convolutional neural network used to extract object embeddings. **C**: Graphical representation of the meta-MDP model of visual search. *F_o_* represents a vector of mean beliefs for a particular object and *1/J_o_* represents a vector of precisions around those beliefs. **D**: Architecture of the deep reinforcement learning agent trained to solve the meta-MDP. A policy and value network each received a set of inputs consisting of object locations, object features, target features and the posterior over which object is the target.

As an initial validation of these two feature spaces, we asked whether the representation can predict fixations in a relatively model-agnostic way. Specifically, we asked whether every time that participants look at an object, the object being fixated is more similar in feature space to the target than to the average of all distractors in the scene (Fig. 3A and B). We found that participants were more likely to look at objects that are similar to the target rather than to an average of all distractors (Fig. 3A and B). Furthermore, the distance between the objects present in a scene and the target was modulated by whether the object was fixated or not (Fig. 3C). In other words, objects that were fixated during search were more likely to be similar to the target than objects that were not fixated. Taken together, these results suggest that the explicit feature space captures some of the underlying perceptual representations that people are using to guide search.

**Fig. 3.**
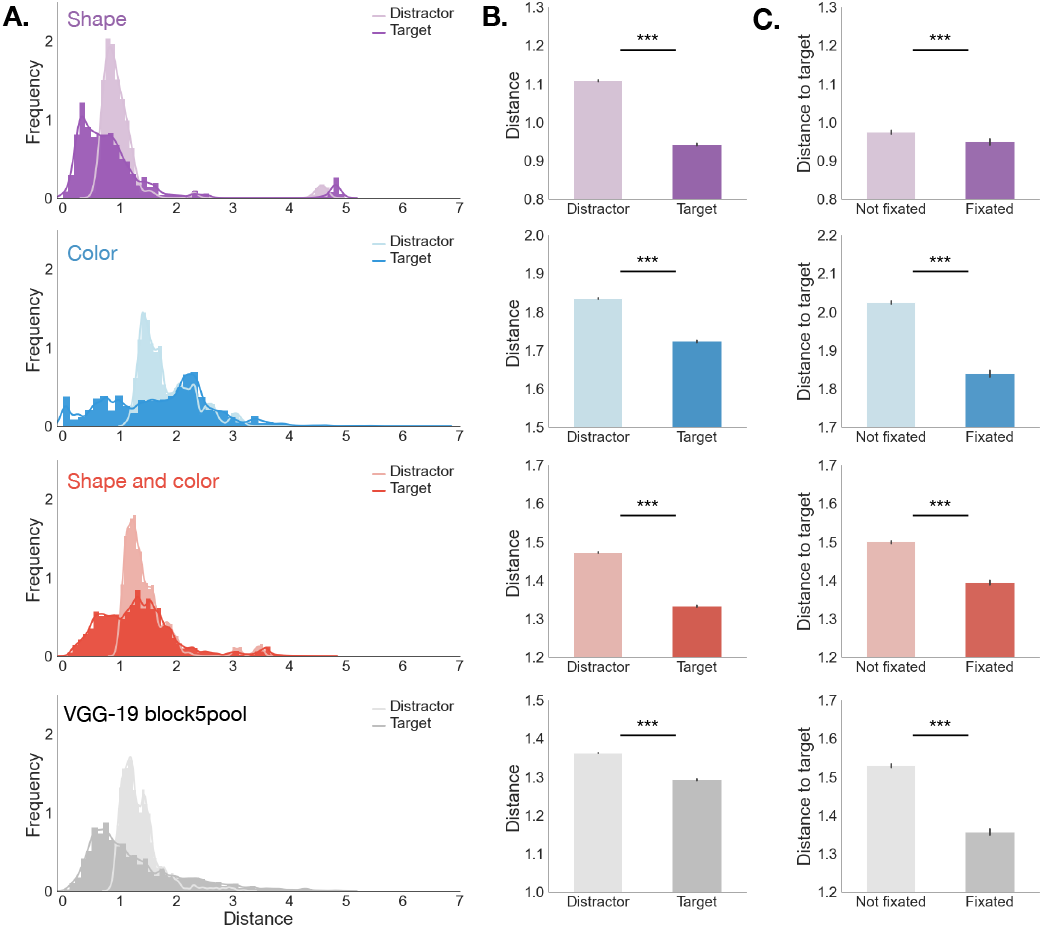
Distance to target in feature space is predictive of gaze. **A**: Distribution of distances between the features of currently fixated object and the features of either the target (darker shade) or the average features of all distractors (lighter shade). This distribution was computed separately for the shape, color, shape + color and VGG-19 feature spaces. **B**: Means and 95% CIs of the distributions in A. **C**: Distance to target for all objects in a scene as a function of whether the object was fixated or not. For the shape, color and shape + color spaces, distance was computed as Euclidean distance. For the VGG-19 embedding, distance was computed pairwise between object as 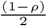, where *ρ* is the Spearman correlation coefficient. ***: *p* < .0001.

Although the previous analysis shows that people are sensitive to both shape and color when choosing where to fixate, people could very well be drawing on a richer representation of the objects. To address this possibility, we also considered a feature space that was learned by a convolutional neural network (CNN). Specifically, we passed a 2D image of each object through the perceptual module of a pretrained VGG-19 network trained on image classification (33) (Fig. 2B, Methods B). Because of their ability to learn highdimensional representations of complex images, CNNs have been proposed as models of the human visual system (34, 35). We focused on the embedding from the last perceptual layer of VGG-19, which was a better predictor of which objects were fixated as a function of target similarity (Methods B, Fig. S1). We reasoned that the representations CNNs learn for image classification might also transfer to visual search in naturalistic environments. Indeed, we found that distance in VGG-19 embedding space predicted both that fixated objects were more similar to the target than to an average distractor, and that fixated objects were in general more similar to the target than objects that were not fixated (Fig. 3, bottom row).

Finally, a challenge for agents learning in multidimensional environments is identifying a low-dimensional state representation that supports behavior (36–38). To compute this representation for our setting, we applied principal component analysis (PCA) to each candidate feature space (shape, color, or VGG-19 embedding). In a subsequent analysis, we verified that the similarity structure of objects was preserved for all three feature spaces, even when retaining a relatively small number of principal components (Fig. S2). Given that fixated and non-fixated objects were discriminable in all three feature spaces (Fig. 3C), we considered all three of these as possible representations in the following analyses.

In addition to validating our choice of representation, the above results indicate that people are using a strategy that fixates objects that are “similar” to the target in some respect. This is consistent with the classic ideal observer models of visual search. To demonstrate this rigorously, we must determine what exactly constitutes an ideal observer in our task.

Representing target and distractor objects in abstract feature spaces is likely necessary to conduct a visual search in a 3D environment. However, it also poses a problem for traditional ideal observer models, which assume that search targets have unstructured image statistics – that is, they can be parameterized as Gaussian and IID (7). Because we are studying the more general case in which these assumptions do not hold (Figs. S3, S4, S5, S6), it is unclear what constitutes “ideal” observing. To answer this question, we first formally define the problem that the searcher is solving.

### B. Defining the search problem

The key intuition behind our analysis is that visual search can be cast as a sequential decision problem (7–12). That is, search consists of a sequence of decisions about where to fixate next, and each such decision takes into account the information gained from previous fixations.

One way to formalize this idea is to frame the problem of computational resource allocation as a *metalevel Markov decision process* (39, 40). Like a standard MDP (41), a meta-MDP consists of a set of states, a set of actions, a transition function, and a reward function. Unlike in a standard MDP, however, the states correspond to beliefs and the actions correspond to computations. The transition function describes how computations update beliefs. Finally, the reward function incentivizes decisions made based on accurate beliefs, but also penalizes computation. Below, we describe a meta-MDP for a general version of visual search (Fig. 2C, see Methods C for the full model).

- **Latent state**: A visual scene is defined as a set of objects at specific locations, with each object represented as a feature vector in a low-dimensional space. We assume that at the beginning of each search episode, the agent knows the location of all objects, but does not know the features of the objects.
- **Belief states**: A belief is a distribution over latent states. For every object, the agent’s current belief is given by two vectors, representing the current mean and precision of that object’s features.
- **Computations**: A computation corresponds to fixating a particular object and sampling information about its visual features. A special computation indicates that search should be terminated (see Reward).
- **Transition function**: The transition function specifies what information is sampled by each computation (the features of objects near the center of gaze) and also how that information is incorporated into beliefs (Bayesian cue combination).
- **Reward**: The agent incurs a cost for each computation. After a computation, it calculates a posterior probability over which object is the target given the current belief state. Search terminates when the maximum of the posterior exceeds a threshold. Upon termination, the agent reports the object most likely to be the target and receives a reward if this is correct and no reward otherwise.

Note that a meta-MDP can be thought of as a special case of a partially observable Markov decision process (POMDP), in which the state does not change and most actions have strictly negative reward. We return to this point in the Discussion.

### C. Learning a resource-rational search policy

By expressing visual search as a meta-MDP over a complex representation of the environment, we can ask, what is the resource-rational (optimal) policy that maximizes performance while minimizing the cost of search?

To address this question, we trained artificial agents to solve the meta-MDP using deep reinforcement learning. We provided artificial agents with the states, computational actions, rewards/costs and transition function that define the dynamics of the environments experienced by human observers, and used proximal policy optimization (PPO) (42) to discover an optimal policy (Fig. 2D, Methods D). Deep reinforcement learning allows us to move beyond scenarios where the optimal policy is analytically computable based on simplifying assumptions.

Using the implicit VGG-19 representation reduced to 6 principal components as a feature space, we trained 10 independent artificial agents and inspected their returns and policy (Fig. 4). We found that all 10 agents achieve asymptotic high returns after about a million training episodes (Fig. 4A). Moreover, agents converged to a policy in which they terminated search whenever the maximum posterior belief exceeded a threshold (Fig. 4B) and consistently fixated the object most likely to be the target (Action 1 in Fig. 4C). We term this policy *Fixate_MAP,* shorthand for fixating the *maximum a posteriori* (MAP) target given the agent’s current belief state. All agents converged to similar returns, which were approximately equal to that of a hard-coded *Fixate_MAP* policy (dotted line in Fig. 4A). We repeated this training procedure with 10 additional agents, this time using the explicit representation defined by shape and color features. While returns were lower on average then for the VGG-19 represen-tation, agents still converged to a policy that approximates *Fixate_MAP* (Fig. S7).

**Fig. 4.**
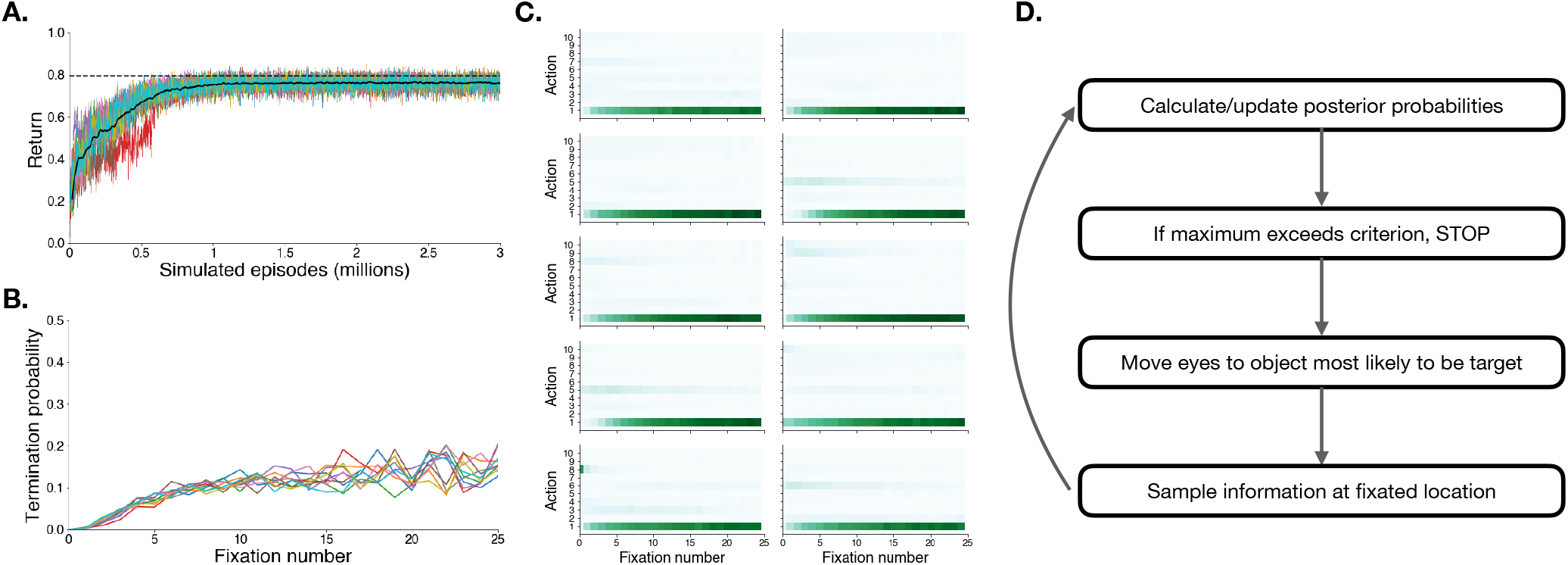
Model training results for VGG-19 feature space. **A**: Learning curves for 10 agents trained to solve the meta-MDP using deep reinforcement learning. Note that all agents converge after about 1 million episodes. The dotted line represents the average return obtained by the *Fixate_MAP* policy depicted in D, over 10,000 simulations with a an optimal stopping criterion. **B**: Probability of terminating search as a function of the number of fixations previously taken, for the 10 different networks. **C**: Histogram of chosen action as a function of fixation number for each of the 10 agents. Actions are ordered by the posterior probability that the fixated object is the target. So Action 1 corresponds to fixating on the object most likely to be the target.**D**: Graphical depiction of the *Fixate_MAP* policy that agents learned.

Notably, the *Fixate_MAP* policy that the agents learned incorporates the previously proposed “ideal observer” model for visual search in simple environments as a special case. For these simple environments, the ideal detector is an inner product of the image with a fixed template (7), which we can interpret in our meta-MDP as a 1D feature space. By using deep reinforcement learning to train agents to solve the more general version of visual search from scratch, we provide a rational basis for this policy in structured, naturalistic settings. All agents converged to the identical *Fixate_MAP* policy, suggesting that *Fixate_MAP* is close to the optimal policy for the meta-MDP. In the analyses that follow, we use *Fixate_MAP* as a computationally tractable approximation of the optimal policy.

### D. Low-dimensional state representations

To verify how sensitive *Fixate_MAP*’s performance is to the choice of representation, we simulated behavior from agents that behave according to *Fixate_MAP* over different versions of the feature space, varying the number of principal components, and optimizing the termination threshold for each resulting representation (Methods E, F). The main finding that emerged from this analysis is that the choice of representation significantly impacted agents’ performance on the task.

First, agents that had access only to explicit shape features achieved lower returns (Fig. S9A and B) and spent a longer time searching (Fig. S10A and B) than agents that had access only to explicit color features. Combining shape and color features gave agents a marginal advantage above and beyond only using color (Fig. S9C and S10C). These findings are not surprising given that, for most objects, the retinal projection is a highly ambiguous signal that maps to 3D shape. Therefore, unlike in research using a fixed set of 2D spatial patterns, shape may not be a useful feature for many visual search tasks characterized by natural 3D scene perception. Notably, agents that learned over the VGG-19 representation performed best (Fig. S9D, Fig. S10D), suggesting that perceptual features useful for image classification also transfer to naturalistic visual search.

Finally, we found diminishing returns in terms of performance as we added more principal components. In other words, the *Fixate_MAP* policy performed well over a relatively low-dimensional state representation. This was true for both average return (Fig. S9) and number of fixations (Fig. S10). Taken together, these findings supported our choice to represent the feature space in the meta-MDP in terms of the first 6 principal components of the VGG-19 embedding.

### E. Predicting human behavior

Our analysis of the policies learned by the meta-MDP provides evidence that Fix-*ate_MAP* is a near-optimal solution to the naturalistic visual search task defined here. But is the *Fixate_MAP* policy consistent with human behavior? To answer this question, we simulated behavior from the policy and compared it to human behavior in the immersive visual search experiment.

#### E.1. Similarities in search performance

First, we qualitatively compared the accuracy (defined as fraction correct) and reaction time distributions obtained in simulation to those observed in the experiment. We simulated the experiment 10 times with a fixed setting of the parameters (cost, threshold, and measurement precision), assuming a linear relationship between number of fixations and reaction time. We found that while the model performs slightly better on average than human participants (Fig. 1C), it is able to reproduce the overall shape of both distributions (Fig. 1C and D).

We also found a small, statistically significant correlation between the average fraction correct by target observed in the human data, and average fraction correct by target simulated by the model (Fig. 1E, Spearman’s *ρ* = .25,*p* = .02). In other words, when the model finds the target, it is more likely to find the same targets that humans do.

Finally, we asked whether the room context modulates accuracy in the same way for participants as it does for model (Fig. 1F). If search is easier for the model in the same contexts in which it is for humans, we might expect a similar ranking in terms of accuracy across rooms. While we did find some consistency – for instance, the “guest bedroom”, “kitchen”, “living room” and “studio” contexts are ranked the same with respect to accuracy for both humans and model – there also are discrepancies. One possible reason is that humans also make use of higher-order semantic priors that we did not explicitly include in the model’s state representation.

#### E.2. Comparing human and model search policies

A key strength of our approach is that we can treat the *Fixate_MAP* policy as a fully generative model of human gaze in naturalistic settings, and ask how well predicted gaze patterns correspond to observed gaze (Fig. 5). To answer this question, we directly simulated gaze behavior from the policy, and compared the resulting gaze traces to those of humans while they performed visual search in virtual reality.

**Fig. 5.**
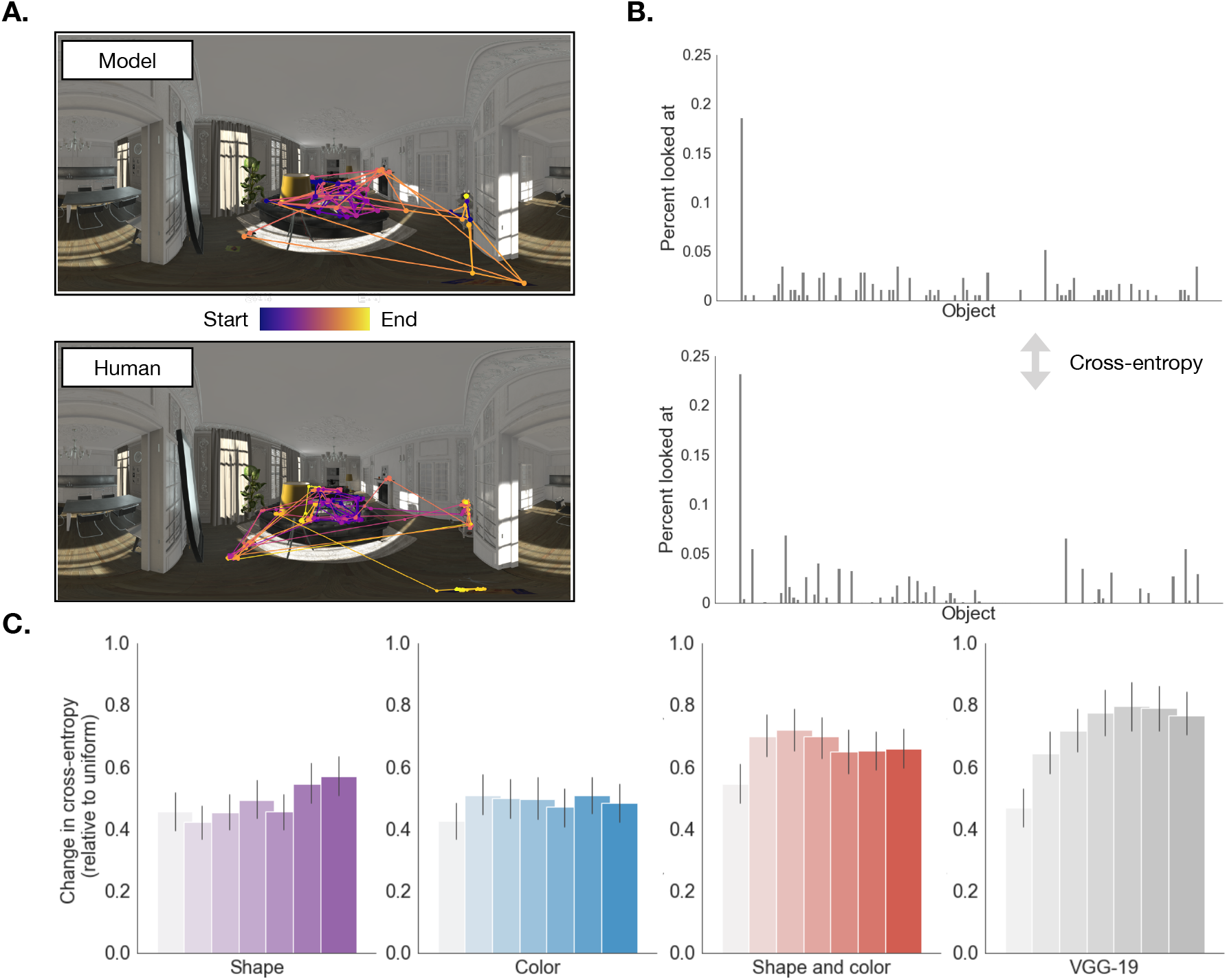
Model predictions. **A, top**: gaze trajectories obtained by simulating the *Fixate_MAP* policy 8 times in one of the living room scenes. **A, bottom**: raw gaze trajectories of 8 participants searching in the same scene, retaining only gaze samples annotated with a task-relevant object. **B, top**: relative looking times to all objects averaged over 20 simulations of *Fixate_MAP* for the scene on the left. **B, bottom**: relative looking times to all objects averaged over participants whose gaze trajectories are shown on the left. Agents were compared based on the cross-entropy match between simulated and empirical relative looking times. **C**: Cross-entropy relative to baseline between simulated and empirical relative looking time distributions, as a function of state representation and number of principal components. Higher bars indicate better fit. Note that for the “Shape and color” representation, the effective number of principal components used was twice each value shown on the horizontal axis. For example, the agent that simulated behavior going into the first red bar had access to one shape and one color feature (two in total).

We found a qualitative match between the temporal structure of participants’ gaze traces and the gaze trajectories simulated by the model. For instance, in the scene shown in Fig. 5A, both the model and humans tended to first look around the couch and coffee table, then at the object on the mantel-piece, then at the objects on the floor, until finding the target on the side table.

To quantitatively compare human and model traces, we calculated the percentage of time the agent looked at each object and computed the cross-entropy between the simulated and observed relatively looking time distributions across all objects in a scene (Fig. 5B, Methods G). Importantly, by making explicit the representation of the environment in the model, we can also investigate the contribution of different features to driving search.

We compared the predictive accuracy of models that generate behavior over each state representation, given different amounts of information quantified as number of principal components (Fig. 5C). A repeated-measures ANOVA revealed that the correspondence between simulated and empirical looking time distributions is statistically significantly different as a function of both feature space (*F*(3, 897) = 87.38,*p* < .0001, *η*^2^ = 0.23) and number of principal components (*F*(6, 1794) = 87.38, *p* < .0001, *η*^2^ = 0.21). The interaction was also statistically significant, suggesting that adding additional principal components improved goodness of fit for some feature spaces and not others (*F*(18, 5382) = 29.29,*p* < .0001,*η*^2^ = 0.09). A post-hoc paired-sample t-test revealed that the top VGG-19 representation outperformed the top shape and color representation (*M*(VGG-19-6PC) = 0.79, *M*(Shape and color-3PC) = 0.72,*t*(299) = 4.5,*p* < .0001). In sum, of the state representations that we tested, VGG-19 tends to perform best at predicting human gaze behavior.

Taken together, our findings suggest a close correspondence between *Fixate_MAP,* a near-optimal solution to the naturalistic visual search problem defined here, and human search policies. Leveraging this policy as a fully generative model of gaze, we find that, given a suitably complex state representation, we can predict where people look while searching for objects in naturalistic environments.

## Discussion

In this paper, we present a resource-rational analysis of human visual search in virtual 3D environments. We cast eye movements as goal-directed actions aimed at gathering information about the visual environment. We specify a meta-level Markov Decision Process (meta-MDP) that optimally trades off long-term utility with the cost of information gathering. To generate candidate state representations for the meta-MDP, we leverage annotations that come by virtue of having the geometric and color representation of virtual scenes, as well as VGG-19, a pre-trained deep convolutional neural network (CNN) optimized for image classification. We then train deep reinforcement learning agents to solve the meta-MDP, and compare samples from the learned policy with human gaze trajectories.

Our results show that agents converge to a near-optimal policy: sequentially fixating the object most likely to be the target, given current beliefs. We provide evidence that people’s eye movements are consistent with this policy. We further find that the policy best matches human gaze behavior when the state is defined by object embeddings encoded in the CNN, suggesting a shared representational basis between naturalistic visual search and image classification. By combining deep neural networks and cognitive models, we show that naturalistic visual search can be understood as resource-rational sequential decision-making over a rich representation of the visual environment.

Several authors have proposed decision-theoretic accounts of goal-directed eye movements during visual search (7–12). In this view, eye movements are generally thought of as sequential actions an agent takes in order to reduce uncertainty about the environment. Visual search can thus be described as a control strategy (policy) designed to detect the location of a target under sensory uncertainty. In related work, some authors have characterized this control policy as the optimal solution to a partially observable Markov decision process (POMDP) (10–12) with the same basic belief updating dynamics as in our meta-MDP model.

Interestingly, for specific environments, the optimal policy can meaningfully deviate from *Fixate_MAP*. For example, in some cases, the first fixation in an optimal sequence of exactly two fixations differs from the optimal single fixation (12). One possibility for why we did not identify such deviations is that the higher dimensionality of our state space prevented the deep reinforcement learning algorithm from discovering a better solution. Another possibility is that the previously observed deviations from *Fixate_MAP* were capturing specific aspects of the experimental setup that do not occur frequently in more naturalistic tasks.

A general limitation of previous work is that policies for solving the sequential information sampling problem have typically been instantiated in simplified environments with statistical structure designed to fit the assumptions of a given algorithm (7). This leaves open the possibility that the efficiency with which humans search in real world environments arises, at least in part, from an ability to detect and represent statistical structure present in real world scenes.

To test this hypothesis, we combine the meta-MDP with CNN scene representations, used as a plausible model of human visual processing (43–45). This view has been validated experimentally by treating CNNs as a general method for extracting high-level structure that supports various visual tasks, and showing that representations learned by CNNs predict human visual judgments at different levels of abstraction (46–48). Most relevant for the case of visual search, different CNN architectures have been widely adopted as predictive models of visual saliency (49–51). However, CNN-based saliency models are not designed to account for the temporal dependencies typical of human eye movements, and usually only predict fixation distributions aggregated over a given input instead of individual scanpaths (52, 53). Our work brings these research threads together in a formal definition of the full naturalistic visual search problem. By explicitly casting gaze behavior as a policy for where to look in a richly structured visual scene, the model we propose simultaneously accounts for some of the representational complexity inherent to the real world (which sequential decision models of search usually sidestep); and for the sequential, goal-directed nature of search (which has been a challenge for CNN models of saliency).

This approach also opens up the possibility for systematically probing which features of the visual environment guide search, given a particular environment and task goal. The results presented here suggest that to find objects, humans search within a feature space common across other visual tasks, such as object recognition and image classification (33, 54). By ablating the model’s state representation and comparing resulting policies with human behavior, we can verify whether a certain feature guides search or not. This logic is broadly consistent with a top-down feature guidance mechanism whereby the goal (finding a particular object) induces a relevant feature space in which perceptual features are processed in parallel, and search proceeds serially at the level of objects (4, 5). We find that the near-optimal policy best predicts human behavior when including all perceptual information encoded in the specific CNN we used (color, shape, texture and higher-order combinations thereof), but compressed into relatively few principal components.

An important limitation of the current work is that the model’s state representation does not consider higher-order information such as scene guidance and semantic knowledge (6, 55). However, there is evidence that humans do employ such inductive biases when searching in real-world scenes. For example, “anchor objects” can be highly informative regarding the likely position of other objects in a scene (56) (e.g., tracking the position of the ball during a game of soccer may provide information about where players are). And different surfaces are more likely to be looked at if they are related to a specific target (e.g., we tend to look for a toaster on a kitchen counter rather than on the fridge). Such information could be captured in the meta-MDP, for example by incorporating object co-occurrence statistics into the state representation.

Our model posits that gaze should be deployed on the basis of both reward and uncertainty, in a manner that minimizes the cost of each fixation (57). However, it is unlikely that humans uniformly collect information across all features of the representation. Depending on the task goal, some features might be more relevant for target detection than others (36, 37, 58, 59). For instance, paying attention to color may be more useful when searching for a piece of clothing than when trying to locate one’s car keys. How such a hierarchy of relevance to visual search may emerge remains to be determined (60). The framework presented here could be extended to incorporate such learning, either by modifying the action space to include sampling at the level of both objects and features, or by letting the cost of sampling different features vary as a function of task goal.

In this paper, we take a first step towards a computational characterization of human goal-directed action selection in fully immersive virtual environments. We show that naturalistic visual search can be understood from the perspective of resource-rational analysis. We provide a computational framework grounded in reinforcement learning for discovering policies that maximize the efficiency of search, while minimizing the cost of gathering information. We instantiate such a policy in a representational space that captures some of the complexity present in naturalistic visual scenes. And we show that human gaze patterns are consistent with this policy. Our findings demonstrate that resource-rational analysis can be leveraged as a predictive model of behavior that generalizes to ecologically valid experimental contexts.

## ACKNOWLEDGEMENTS

We are grateful to Ruta Desai, Kara Emery and Todd Gureckis for insightful discussions on the work presented in this manuscript.

## Methods and supplementary figures

### A. Immersive visual search task

Classic studies of visual search are designed to test the human ability to use specific features to find objects (61). While these studies have identified some features of the environment that drive eye movements, such stimuli do not reflect the multidimensional structure of real-world environments. To overcome this limitation, we conducted a study of visual search in virtual reality (VR). 26 participants viewed VR scenes generated with the Unity game engine through a head-mounted display (HTC Vive VR), equipped with a Tobii Pro VR eye tracker (sampling frequency: 120Hz), while holding a handheld controller.

Participants provided informed consent consistent with the Declaration of Helsinki. Experimental sessions lasted approximately 60 minutes, for which we compensated participants $50. We excluded data from 5 participants for data quality issues, yielding a dataset of 21 participants.

Each participant performed 300 trials of visual search for a target in a cluttered room with 55-112 distractors and under an 8 second deadline. A trial was defined by a combination of one of 6 possible rooms (kitchen, living room, bathroom, study or one of two bedrooms), one of 5 pre-determined viewpoints, and one of 10 pre-generated target/distractor sets. Each participant experienced the set of 300 unique trials this procedure generated but in a different random order. In some of the trials, visual recommendations, in the form of transparent blue blobs, were presented. For each participant, assistance was provided on 100 trials. Data from those trials are not reported here.

Each trial began by showing the target object for 4 seconds in an empty, white scene. The participant was then immersed into the room from a perspective selected *a priori.* At any time during the 8-second search period, the participant could report having found the target by pointing and clicking with a handheld controller. If the participant did not find the target with the 8-second period, the trial ended. We determined the identity, size, location and orientation of the target object and distractor objects with a number of constraints: (1) each object rested on a stable surface (e.g., floor, table, etc); (2) none of the distractors had the same object identity as the target, or are too similar; (3) at least 50% of the target was visible from the participant’s viewpoint; (4) the visible area of the target is at least 3 degrees of visual angle squared.

### B. Feature extraction

#### B.1. Explicit method

Our explicit feature extraction approach draws from work in visual cognition showing that eye movements are guided by objects and object shape, color and texture (5). We quantified shape as the D2 distribution over the 3D mesh of each object (32) (Fig. 2A, left). And we quantified color by converting the 2D texture of each object to the CIE L*a*b* color space and extracting the A (green-red) and B (blue-green) channels (Fig. 2A, right).

The D2 shape distribution and CIE L*a*b* color representation both yield high-dimensional representations of the shape and color of each object, constrained by the number of times pairs of points are sampled from the mesh (in the case of D2) and the pixel size of the texture (in the case of CIE L*a*b*).

To compute the shape distribution, we randomly sampled 500,000 pairs of points on the mesh surface and computed the Euclidean distance between each pair. We further reduced this distribution to a histogram with 100 bins, applying PCA to a shape matrix consisting of *N*_objects_ × *N*_bins_. For converting color texture to CIE L*a*b*, we resized each texture image to 500 × 500 pixels, and concatenated the A and B channels. To preserve spatial color information, we opted not to histogram this representation, applying PCA directly to a color matrix of dimensionality *N*_objects_ × *N*_pixels_.

Due to its ease of computation and robustness in the presence of rotations, translations and other shape perturbations, the D2 distribution has been widely used in the computer graphics and computer vision literature as a metric for discriminating between shapes of different classes (32). Importantly, data from human similarity judgments suggests the D2 distribution captures some aspects of human shape perception (62).

The CIE L*a*b* color space describes all the colors visible to the human eye, and can thus serve as a metric space for color similarity that roughly matches that of human color perception. Because color was extracted from 2D texture files, this representation is likely to also include some textural elements such as smoothness or graininess.

#### B.2. Implicit method

Recent work has shown that CNNs capture some aspects of the human visual architecture and processing (43). 2D images of objects can be represented as points in multidimensional feature spaces learned by CNNs (known as “embeddings”). These embeddings can then be used downstream to support different tasks such as categorization, typicality judgments or visual production (34, 47, 63). We used a standard pre-trained Keras implementation of the VGG-19 architecture (33) to extract embeddings across different depths of the network (Fig. 2B).

Specifically, we passed a 224 × 224 pixel image of each object viewed from a canonical viewpoint through a Keras implementation of the pre-trained VGG-19 architecture. Running the model forward, we obtained a multidimensional embedding of all objects in the experiment. How predictive this embedding was of gaze depended on which layer of the network we used. VGG-19 consists of two modules: a perceptual module that alternates blocks of convolutions with pooling layers; and a classification module consisting of two fully connected layers and a softmax layer. We analyzed feature activations in the second-to-last fully-connected layer (FC-4096, dimensionality 1 × 4096), as well as the last pooling layer (Max-Pool 5, dimensionality 7 × 7 × 512). For the latter, we averaged and flattened the 512 spatial maps, yielding an embedding with 49 dimensions for each object.

Qualitatively, the embedding computed from the perceptual module was a better predictor of gaze than the embedding computed from the classification module (Fig. S1A). However, distance computed from both embeddings was predictive both of whether a particular fixation would fall on an object that is more similar to the target than an average distractor (Fig. S1B), and of whether fixated objects were more similar to the target (Fig. S1C).

Quantitatively, we found a weak but significant difference between layers in the strength of both of these relationships. The difference in mean distance between the object being fixated and target, and mean distance between the object being fixated and an average distractor, was slightly higher for the classification embedding (*M* = 0.076) than for the perceptual embedding (*M* = 0.068); a two-way ANOVA revealed a significant interaction between layer and object type (*F*(3, 1321420) = 8.65,*p* < .01). However, the difference in mean distance between target and fixated vs. not-fixated objects was higher for the perceptual embedding (*M* = 0.173) than for the classification embedding (*M* = 0.114), with a two-way ANOVA revealing a significant interaction with layer (*F*(3, 343332) = 74.14,*p* < .0001). Since the perceptual embedding was a better predictor of which objects were fixated as a function of target similarity, we used this embedding in subsequent modeling analyses.

#### B.3. PCA

To further reduce the dimensionality of our feature space, we separately applied PCA to each feature type and computed low-dimensional projections in the space defined by the principal components. For the shape and color features, PCA was applied over matrices in which rows indexed objects, and columns indexed bins of either the shape histogram or raw pixel-level color information associated with each object. For VGG-19 features, columns indexed features of each object’s embedding obtained from the Max-Pool 5 layer. Across all features, we computed the principal components needed to explain 95% of the variance in feature space.

### C. Full meta-MDP specification

A meta-MDP consists of a latent state, beliefs over the latent state, actions, a transition function and a reward function. The full meta-MDP for visual search can be specified as follows:

- **Latent state**: The latent (unknown) state is represented as a matrix *A* of dimensionality *N_o_* × *Nf* such that the value of feature *f* for object *o* is *A_of_*, where *N_o_* is the number of objects and *N_f_* is the number of features in a particular representational space. This matrix is assumed to be a property of the environment, but it is not known to the agent when search begins.
- **Beliefs**: The agent represents the environment as beliefs over features in *A*. For tractability, we assume independent Gaussian beliefs. Beliefs can thus be represented with two mean and precision matrices of dimensionality *N_o_* × *N_f_*, *F* and *J*, such that 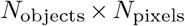. Here, *F_of_* represents the mean belief for a particular feature *f* of object *o*, and 1 */J_of_* represents the precision around that belief.
- **Computations**: A computation corresponds to looking at the center of object *o*.
- **Actions**: An action corresponds to performing a computation.
- **Transition**: Formally, a computation takes a measurement *X* of the features of objects near the center of gaze, and incorporates that measurement into the belief using Bayesian cue combination. To specify this transition:

1. Compute an object attentional mask *g_o_,* in which the attention paid to all objects in the scene exponentially decreases with their distance to *o*.
2. Compute the measurement precision *J*_meas_ = *g_o_1^T^* such that the features of objects near the center of gaze have the highest precision.
3. Independently sample a measurement from the true latent state:

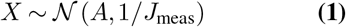
4. Perform the Bayesian update

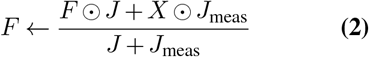

and *J* ← *J* + *J*_meas_, where Θ represents the element-wise product.
- **Reward**: The agent incurs a cost –*c* for each computation. When the agent terminates computation, it calculates a posterior over which object *o* is the target given its beliefs,

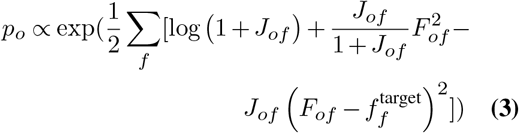 This calculation assumes that the true values of object features are all mutually independent and Gaussian with mean 0 and variance 1. The agent then selects argmax[*p_o_*] as its target report and receives a final reward of *R* = 1 if this is correct and *R* = 0 otherwise.
- **Termination**: We denote ⊥ as a special computation for which search terminates whenever max[*p_o_*] exceeds a threshold.

### D. Solving the meta-MDP

In order to identify the optimal policy for the meta-MDP, we trained artificial neural network agents to maximize the expected cumulative rewards in an ensemble of visual search environments. For each of the 300 scenes presented in the human task, we generated an artificial environment in which the target was the same as in the human experiment. We also created additional environments by considering the same scene, but with a different object assigned to be the target. We only generated such scenes if that object was unique among the object set, resulting in an ensemble of 7022 environments.

Artificial environments were generated from the first 6 principal components of the feature representations for all objects in the scene. To ensure that each environment has the same number of objects, which is necessary for our network architecture, we added ‘phantom objects, to each environment with less than the maximum number of objects (113). For these phantom objects, we set the true feature values to zero. We made one additional modification to the state and action space. When passing the belief state to the preprocessing layer, we re-ordered objects by rank order according to the posterior, and we applied the same re-mapping for the action space. In other words, the first rows of the belief state, and the first entry of the action space, always correspond to the object that is most likely to be the target according to the belief state; and action 1 corresponds to fixating on that object.

We implemented and trained the artificial agents in tensorflow (64) with the `tf-agents library` (65). We used proximal policy optimization (PPO) (42) to define the loss function, and trained using Adam (66) with exponential learning rate decay.

Since PPO is an actor-critic method, we specified two neural networks which only differed in the output layer (Fig. 2D). Both networks consist of a preprocessing layer which represents inputs (see below), then a series of 3 densely connected layers of 113 units each, followed by an output layer (a single value for the critic, an action distribution layer for the actor).

The preprocessing layer first represents a belief state in the meta-MDP as a list of 4 tensors

- *F*, a 113 × 6 matrix of the mean feature values.
- *J*, a 113 × 6 matrix with the corresponding precisions.
- *f*_target_, a 1 × 6 vector with the true values of the target object’s features.
- *x_o_, a* 113 × 2 matrix with the horizontal and vertical coordinates of each object, normalized so that the screen maps to the interval [–1, 1]. For phantom objects, we set *x_o_* to (0, 0), the screen center.
- *p_o_*,a 113 × 1 vector with the output of Equation 3. We set *p_o_* to 0 for phantom objects, then re-normalize to ensure the sum of *p_o_* over non-phantom objects equals 1. Although *p_o_* is a deterministic function of *F*, *J* and *f*_target_ and therefore provides no additional information to the agent, we include it in the state representation to accelerate learning.

The preprocessor then passes each of these tensors through a single dense layer with 113 units and concatenates the outputs into a single tensor which represents the belief state.

### E. Threshold optimization

The *Fixate_MAP* policy terminates search when any item is deemed sufficiently likely to be the target. Formally, the policy executes the ⊥action when max[*p_o_*] > *θ*, where the threshold *θ* is a free parameter. We set this parameter to maximize the expected total reward attained per trial. Concretely we defined an objective function that approximates the expected reward as the average reward in 10,000 simulated trials, and optimized this function with a grid search over the range (0.5, 0.999) in logit space (i.e., smaller distances between values near 1).

### F. Ablation study

To determine how sensitive reinforcement learning agents are to the choice of representation, we performed an ablation study in which we simulated behavior from the *Fixate_MAP* policy over different state representations, using a threshold optimized for the specific state representation (Fig. S8). Specifically, we simulated behavior from the agent over shape features only, color features only, shape and color features and VGG-19 features, varying the number of principal components that the agent had access to. And we computed the average return and number of fixations that *Fixate_MAP* agent obtained given these different state representations.

### G. Comparing human and agent policies

To analyze gaze trajectories, we first transformed the raw gaze sample coordinates to the pixel space of 360 degree equirectangular projections from the pre-specified viewpoints in each scene (67) (Fig. 1B). This transformation is necessary in order to simultaneously take into account participants’ head and eye movements.

We did not apply other processing or smoothing to gaze data, and we did not segment trajectories into fixations and saccades, since saccade detection algorithms are developed for head-fixed participants viewing 2D screens and may not readily extend to free viewing in 3D virtual reality environments.

We then annotated each raw gaze sample with the label of the object at the center of gaze. We extracted this object label by recording the object mesh collider hit by a ray in the Unity environment in the direction of gaze. We found that about 50% of gaze samples landed either on task-relevant objects (i.e., the target or a distractor), suggesting that objects are a strong cue for gaze, consistent with previous findings (68). In this paper, we restrict analysis to those gaze samples which hit task-relevant objects.

To compare the policies of the trained agent with those of human participants, we simulated 20 fixation trajectories per scene (Fig. 5). This simulation assumed a fixed cost parameter and a threshold optimized for the specific state representation (Methods E). We then computed the proportion of time (in units of raw fixations) that the agent looked at each object for a given scene. To compare each resulting scene-wise probability distribution with the empirical distribution, we used cross-entropy, fitting a lapse rate for each individual participant. We report this cross-entropy metric averaged across scenes and participants, and subtracted from a baseline model which assumes uniform relative looking times across all objects (i.e. maximal cross-entropy). The higher this difference, the more closely the distribution of objects fixated by the model matches the distribution of objects fixated by participants.

**Fig. S1.**
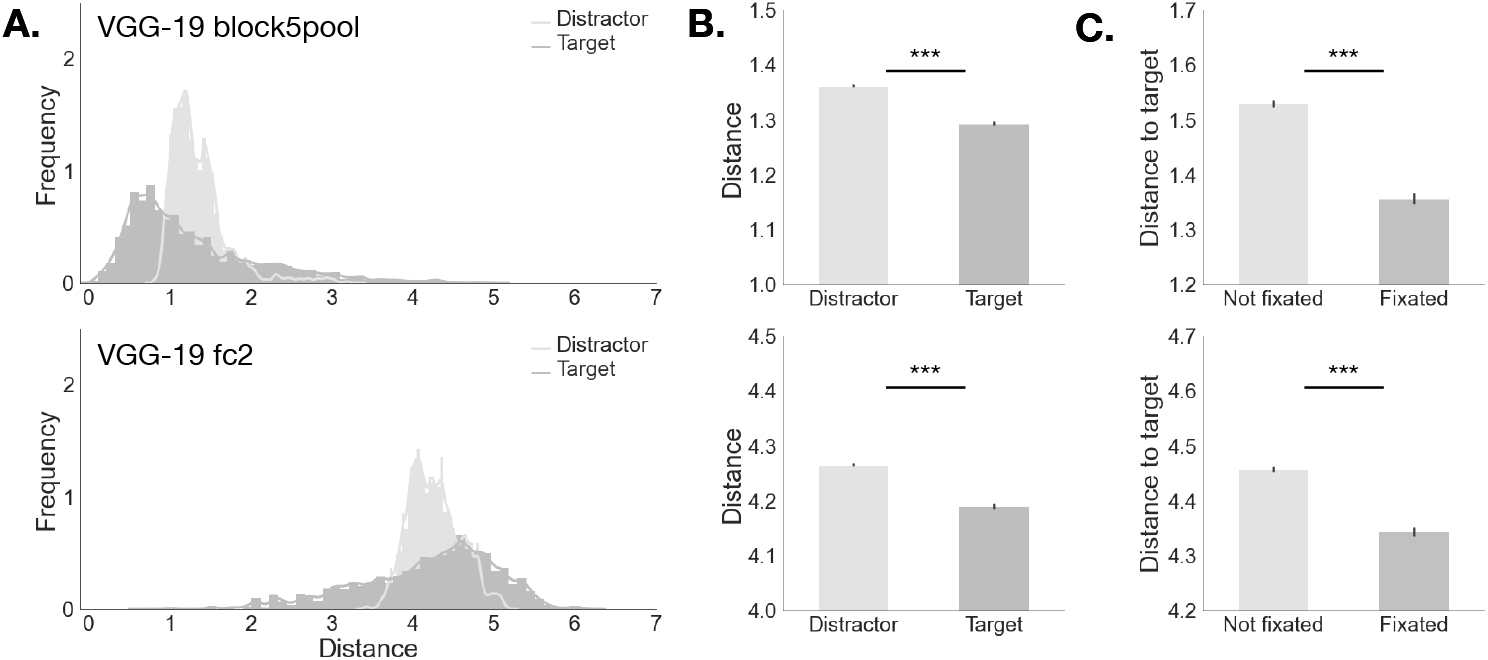
Distance to target in feature space is predictive of gaze. **A**: Distribution of distances between the features of currently fixated object and the features of either the target (darker shade) or the average features of all distractors (lighter shade). Shown here are the distributions computed from the VGG-19 embedding, using either the last perceptual layer of the network (top) or the second-to-last classification layer (bottom). **B**: Means and 95% CIs of the distributions in A. **C**: Distance to target for all objects in a scene as a function of whether the object was fixated or not. Distance was computed pairwise between object as 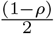, where *ρ* is the Spearman correlation coefficient. ***: *p* < .0001.

**Fig. S2.**
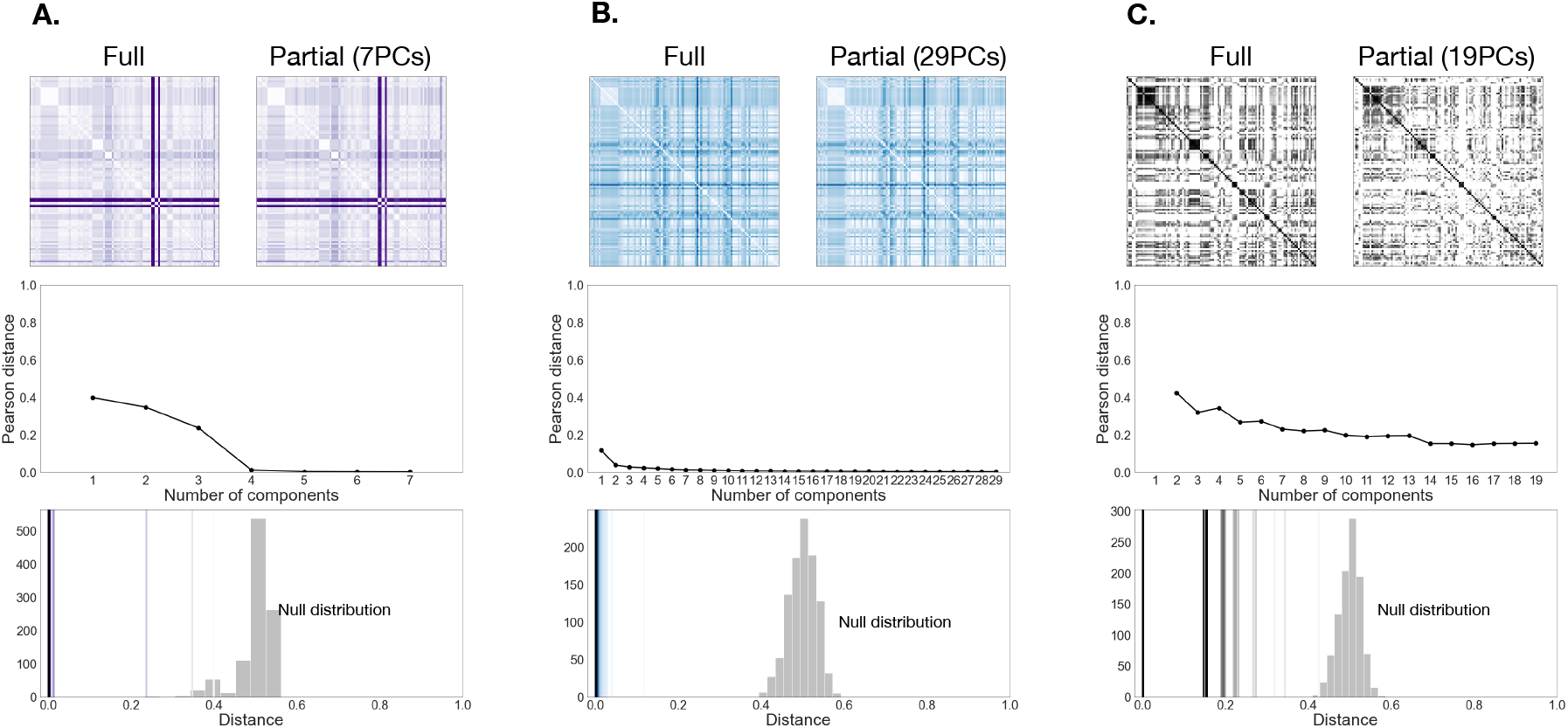
PCA validation. To validate the choice of PCA as a dimensionality reduction step in our model, we generated representational dissimilarity matrices (RDMs) for all the objects in used in the experiment (69, 70). For each object we computed the pairwise Euclidean distance between shape and color vectors, and the pairwise Spearman distance between VGG-19 embeddings. We used used either the full representation (RDM-full), or a partial representation based on a restricted number of principal components (RDM1, RDM2, etc). An example comparison between the full RDM and a partial RDM is shown in the top row for shape (A), color (B) and the VGG-19 embedding (C) respectively. We found that for all representational spaces, the distance between full and partial RDMs decreases as the number of components used increases, middle row). We then performed a permutation test to obtain a null distribution for the Pearson distance between the intact and randomly permuted full RDM. This distribution provides an estimate of the upper bound on the Pearson distance we might expect between different RDMs. Taking the distance of the full RDM to itself as a lower bound, we computed and plotted the distance between RDM-full, RDM1, RDM2, etc. We found that similarity is largely preserved for low-dimensional projections in the space of principal components. For shape, color and the VGG-19 embedding, all RDMs were well outside the null distribution obtained via permutation testing (bottom row). In other words, PCA preserves the representational structure of each candidate feature space.

**Fig. S3.**
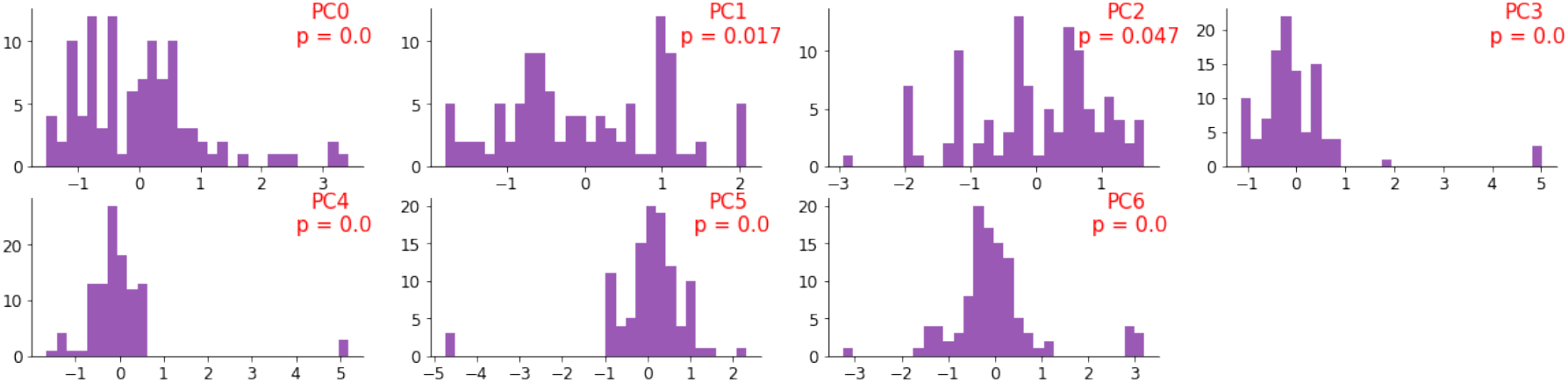
Histogram of shape features, annotated with p-values for D’Agostino and Pearson’s normality test. Red denotes features for which this test was statistically significant (that is, we can reject the null hypothesis that features were normally distributed).

**Fig. S4.**
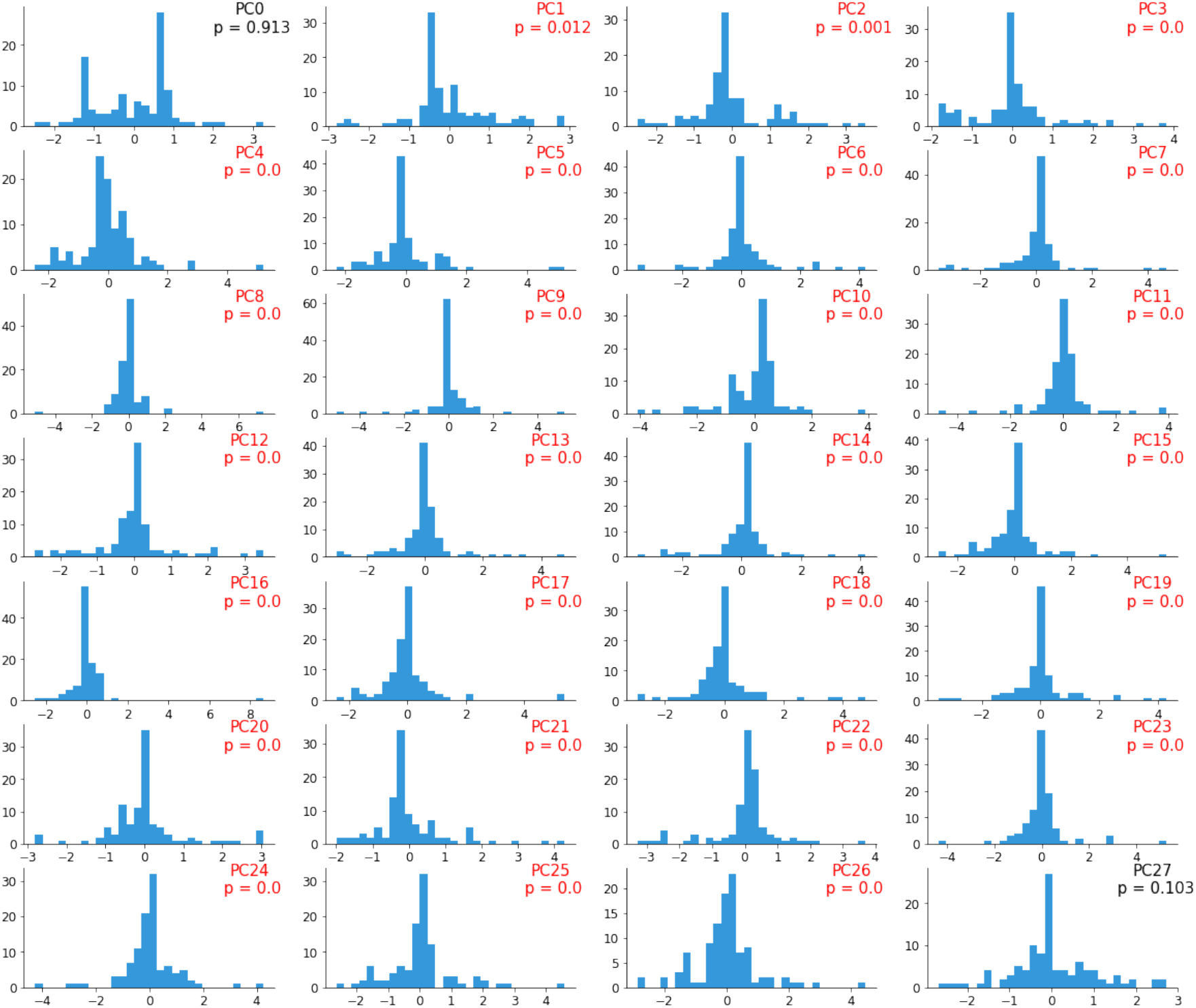
Histogram of color features, annotated with p-values for D’Agostino and Pearson’s normality test. Red denotes features for which this test was statistically significant (that is, we can reject the null hypothesis that features were normally distributed).

**Fig. S5.**
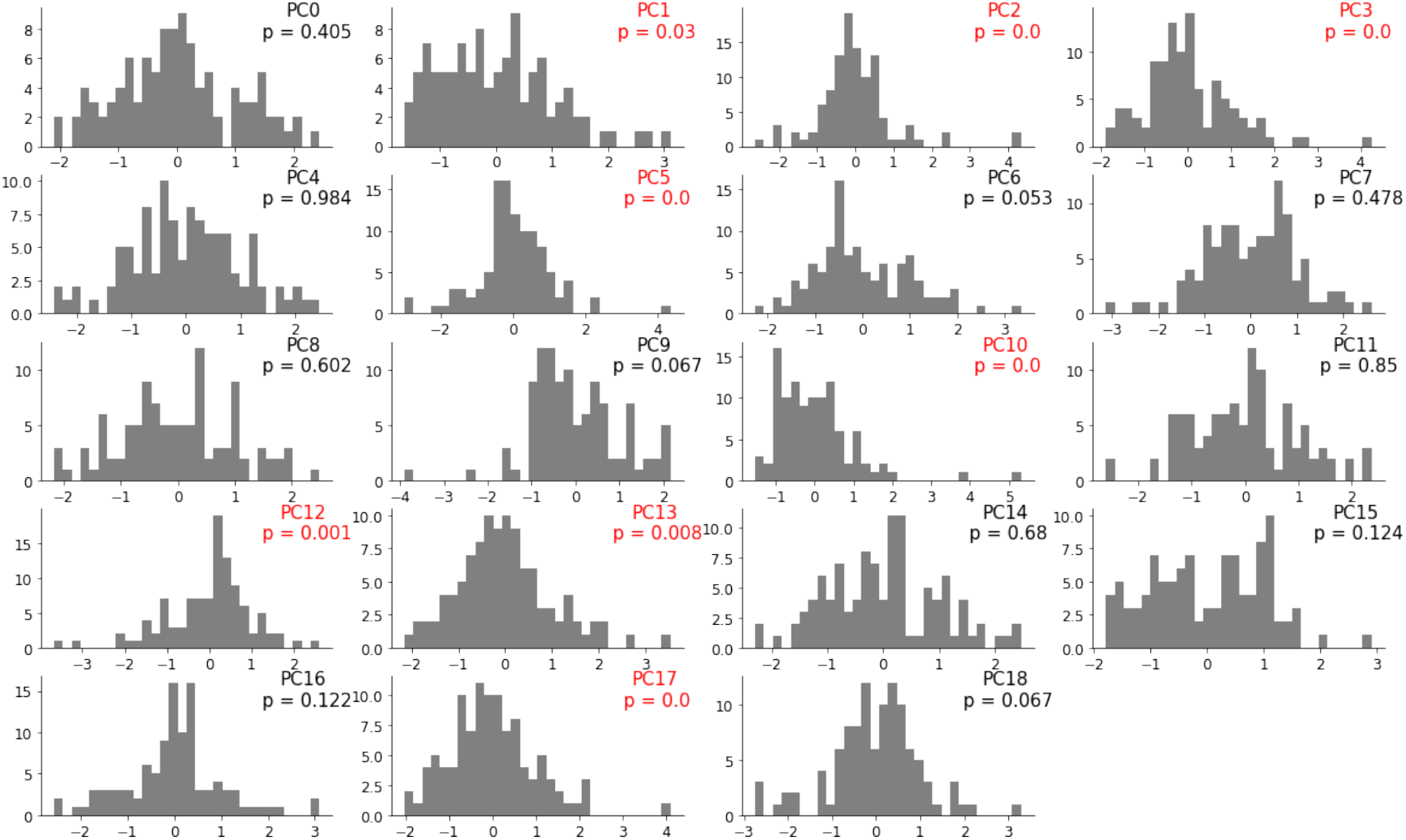
Histogram of VGG-19 features, annotated with p-values for D’Agostino and Pearson’s normality test. Red denotes features for which this test was statistically significant (that is, we can reject the null hypothesis that features were normally distributed).

**Fig. S6.**
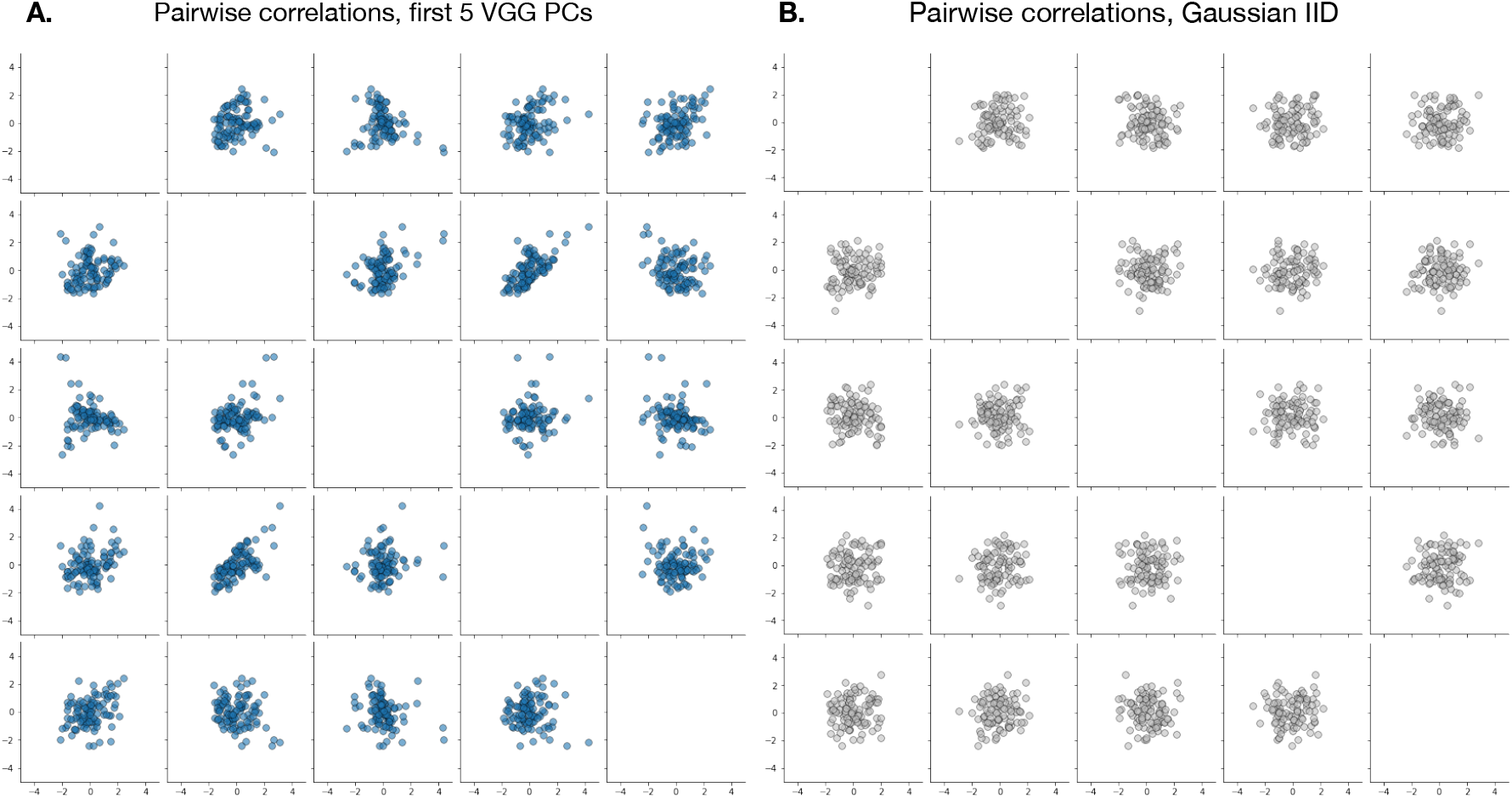
Comparison of pairwise correlations between VGG-19 features (**A**) and randomly generated Gaussian IID features (**B**). If features were Gaussian IID, then we would expect the covariance of any two features to be close to 0.

**Fig. S7.**
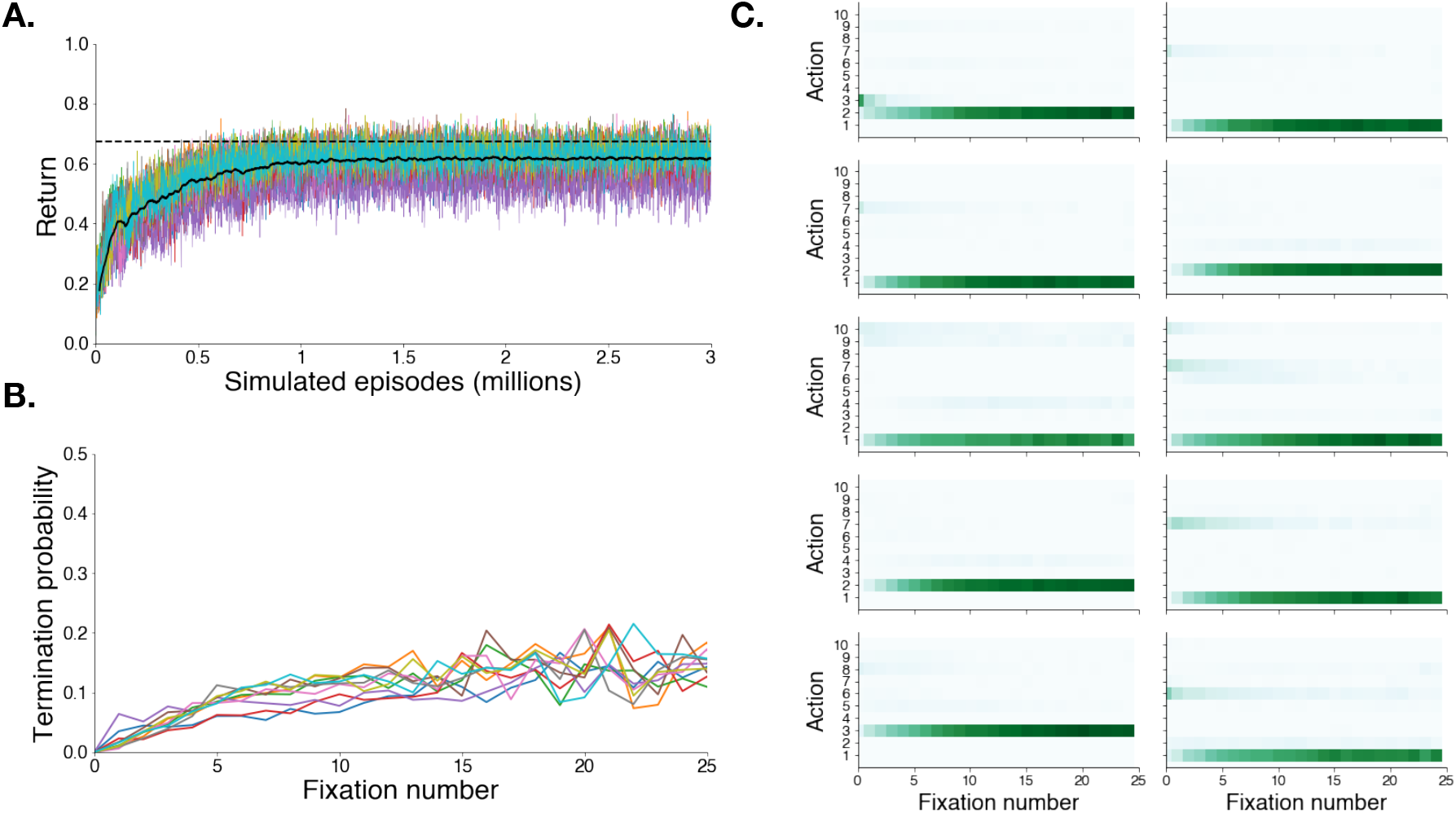
Model training results for shape and color feature space. **A**: Learning curves for 10 agents trained to solve the meta-MDP using deep reinforcement learning. As for the VGG-19 representation, all agents converge after about 1 million episodes. The dotted line represents the average return obtained by the *Fixate_MAP* policy (see Results section). **B**: Probability of terminating search as a function of the number of fixations previously taken, for the 10 different networks. **C**: Histogram of chosen action as a function of fixation number for each of the 10 agents. Actions are ordered by the posterior probability that the fixated object is the target. So Action 1 corresponds to fixating on the object most likely to be the target.

**Fig. S8.**
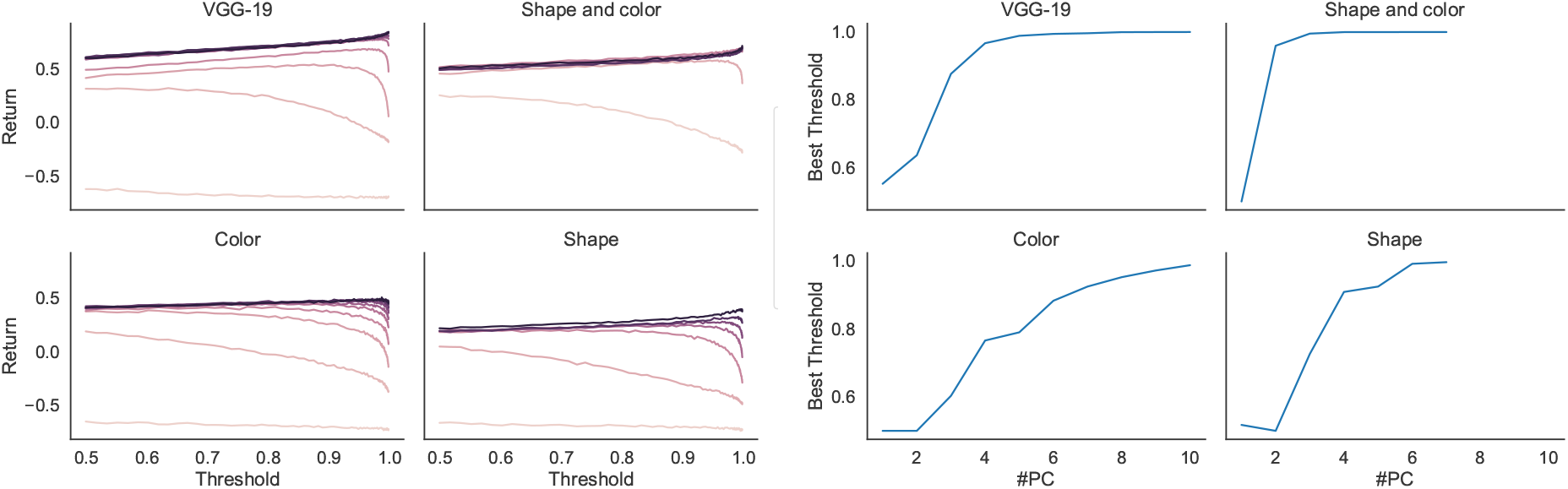
Threshold optimization. Left: the return attained by the *Fixate_MAP* policy with different feature spaces and threshold values. Right: the best-performing threshold for each feature space.

**Fig. S9.**
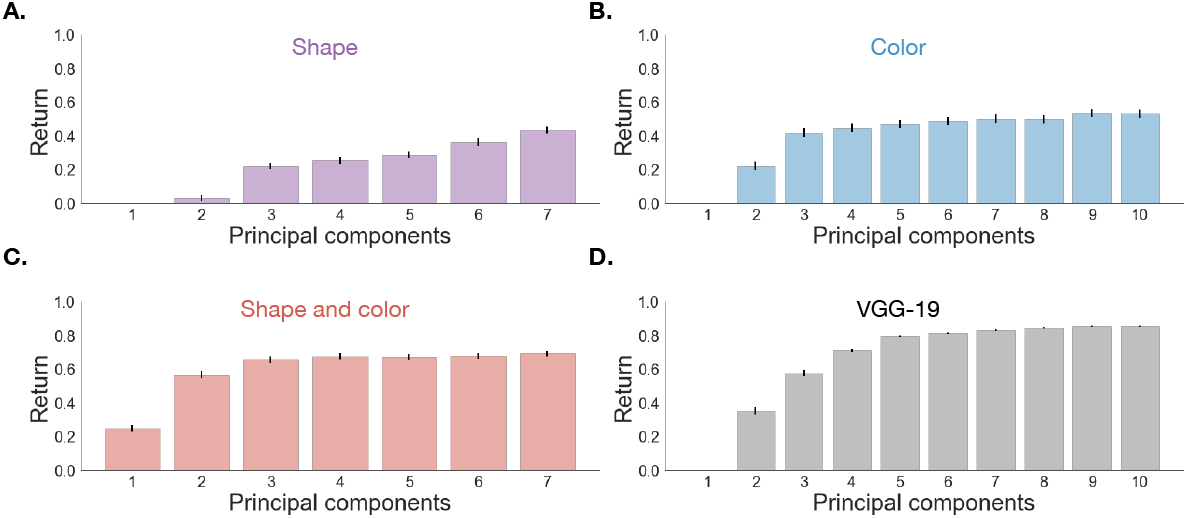
Ablation study return. Average return over many simulations of the *Fixate_MAP* policy, as a function of state representation and number of principal components.

**Fig. S10.**
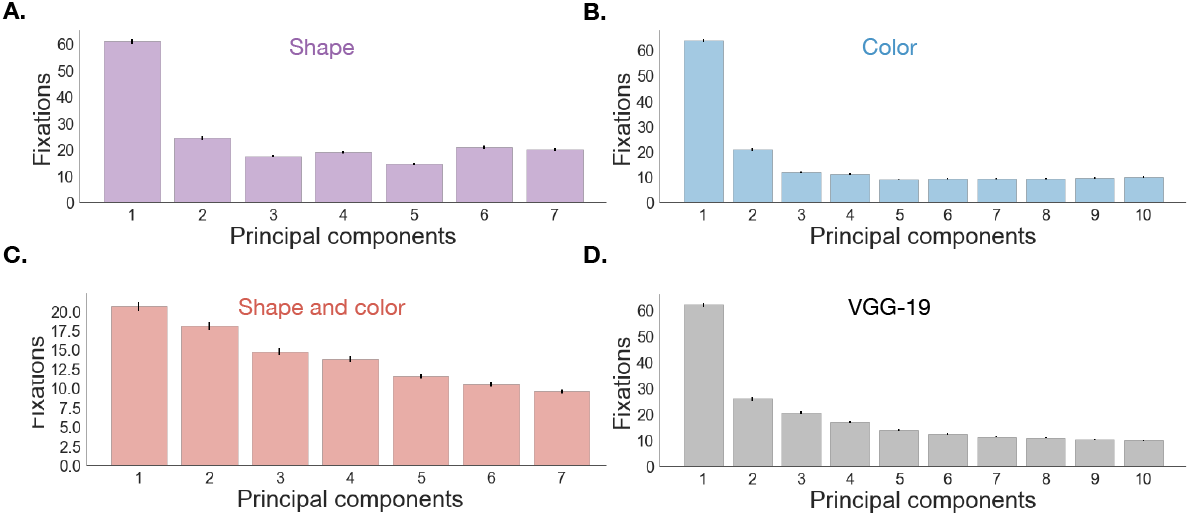
Ablation study fixations. Average number of fixations over many simulations of the *Fixate_MAP* policy, as a function of state representation and number of principal components.

